# Unrevealing water and carbon relations during and after heat and hot drought stress in *Pinus sylvestris*

**DOI:** 10.1101/2021.06.29.450316

**Authors:** Romy Rehschuh, Nadine K. Ruehr

**Affiliations:** Karlsruhe Institute of Technology KIT, Institute of Meteorology and Climate Research, 82467 Garmisch-Partenkirchen, Germany

**Author notes:** Correspondence: Romy Rehschuh.

**Keywords:** electrolyte leakage, heat and drought stress, leaf hydraulic conductance, photosynthesis, root respiration, recovery, Scots pine, stem hydraulic conductivity, transpiration

## Abstract

Forests are increasingly affected by heatwaves, often co-occurring with drought, with consequences for water and carbon (C) cycling. However, our ability to project the resilience of trees to an intensification of hot droughts remains limited. Here, we used single tree cuvettes (n=18) allowing us to investigate transpiration (*E*), net assimilation (*A*_net_), root respiration (*R*_root_) and stem diameter change in Scots pine seedlings during gradually intensifying heat or drought-heat stress (max. 42°C), and post-stress. Further, we assessed indicators of stress impacts and recovery capacities.

Under heat stress, well-watered seedlings prevented overheating of leaves effectively via increased *E*, while under drought-heat leaf temperatures increased to 46°C. However, leaf electrolyte leakage was negligible, but F’_v_/F’_m_ declined alongside *A*_net_ moderately in heat but strongly in drought-heat seedlings, in which respiration exceeded C uptake. Further, the decrease of needle water potential (*ψ*_Needle_) to −2.7 MPa and relative needle water content (RWC_Needle_) under drought-heat reflected a decline of leaf hydraulic conductance (*K*_Leaf_) by 90% and stem hydraulic conductivity (*K*_S_) by 25%. Alongside, we observed pronounced stem diameter shrinkage.

Heat stress alone resulted in low functional impairment and all measured parameters recovered fast. In contrast, larger impacts following combined heat and drought led to the incomplete recovery of *K*_Leaf_ and *K*_S_. Despite *A*_net_ tended to be reduced albeit F’_v_/F’_m_ had recovered, the seedlings’ net C balance reached control values 2 d after stress release and stem growth rates exceeded control rates in the 2^nd^ week post-stress. This indicates that a new equilibrium of C uptake and release was maintained at the tree level, slowly supporting regaining of stress-induced losses.

In summary, we highlight that under moderate heatwaves with low functional impairment, recovery is fast in Scots pine, while in combination with drought hydraulic and thermal stress are intensified, resulting in functional damage and delayed recovery processes. The incomplete recovery of hydraulic conductance indicates limited water transport capacities that could become critical under repeated heat events.

## Introduction

In the past two decades, extreme summer heatwaves and drought periods, combined with high vapor pressure deficit (VPD; Grossiord et al., 2020), have been increasing in Central Europe (Ciais et al., 2005; Ionita et al., 2017; IPCC, 2018). This has increased the pressure on forests, thus reducing primary productivity and enhancing hydraulic constraints, which might lead to decreased vitality and/or tree death in the long-term (Anderegg et al., 2019; McDowell et al., 2020; Schuldt et al., 2020; Arend et al., 2021). Global climate change is forecasted to continue and includes -besides the ecological aspect- the decline of economically important tree species (Hanewinkel et al., 2013) such as Scots pine (*Pinus sylvestris* L.). Although considered a rather drought-resistant species in Central Europe (Ellenberg and Leuschner, 2010), heat waves and drought spells are increasingly limiting its survival, causing c. 40% of the European forest mortality events (Allen et al., 2010).

Studies on the resistance of conifers to thermal stress are limited (Escandón et al., 2016). Further, few experimental studies have focused on the responses of trees to a combination of heat and drought stress (e.g. Zhao et al., 2013; Bauweraerts et al., 2013, 2014; Birami et al., 2020; Kumarathunge et al., 2020), and even less have addressed subsequent recovery trajectories (Ruehr et al., 2016, 2019; Birami et al., 2018). In general, our understanding of tree physiological recovery from combined heat waves and drought stress is very limited (Ruehr et al., 2019), thus hindering our ability of forecasting tree responses to climatic extremes. Recently, Ruehr et al. (2019) suggested that recovery trajectories depend on stress impact, indicating that physiological recovery will be slower if stress causes functional impairment and/or damage. Hence, to understand recovery trajectories, the physiological stress impacts need to be quantified properly.

High temperatures affect the water and carbon (C) relations of trees. C uptake via photosynthesis is closely linked to stomatal conductance (*g*_s_) and water loss via transpiration (*E*). During heat, increasing *E* can cool leaves if soil water is not limiting, while stomata have been found to remain at least partially open (Urban et al., 2017b; De Kauwe et al., 2018; Drake et al., 2018). Drought stress, however, exacerbates during heat waves as the evaporative demand increases, forcing water loss from trees. Drought-induced stomatal closure typically delays the decrease of tree-internal water potentials to critical values, but impedes transpirational leaf cooling, thus leading to higher leaf temperatures (Scherrer et al., 2011; Birami et al., 2018; Drake et al., 2018). As a result of high temperatures, photosynthetic processes might be affected by decreased activity of Rubisco activase and photosystem II (PSII), reduced CO_2_ solubility, and inhibited electron transport (Schrader et al., 2004; Smith and Stitt, 2007; Sage et al., 2008; von Caemmerer and Evans, 2015; Birami et al., 2018). Excessive leaf temperatures may further lead to cell membrane damages in leaves, typically observed via electrolyte leakage (Saelim and Zwiazek, 2000; Correia et al., 2013; Escandón et al., 2016). This often involves the buildup of reactive oxygen species (Demidchik et al., 2014; Larkindale and Knight, 2002), which have been shown to even further impair cell membranes and proteins (O’Kane et al., 1996). The impairment of extra-xylary leave tissues, resulting in leaf hydraulic conductance (*K*_Leaf_) decline, has been found manifold in response to drought, with leaf xylem embolism formation at higher levels of dehydration (e.g. Cochard et al., 2004; Brodribb and Cochard, 2009; Johnson et al., 2009; Skelton et al., 2017). Less information on hydraulic responses to high temperature stress exist and responses of *K*_Leaf_ range from no effect to extreme heat stress (Drake et al., 2018) to increases under modest temperature increments related to changes in the viscosity of water and the symplastic conductance (Sellin and Kupper, 2007; Way et al., 2013).

Besides direct impacts of high temperature and drought on assimilation (*A*_net_), also respiratory processes are affected (Gauthier et al., 2014; Birami et al., 2020). Root respiration (*R*_root_) provides energy for maintenance and growth, necessary for root water and ion uptake (Atkin et al., 2000). However, under heat and hot drought, *R*_root_ has been found to decline, with reduced C availability being one of the reasons (Birami et al., 2020). Yet, C is also necessary to sustain aboveground growth. So far, the knowledge of growth processes in response to heat waves remains elusive, and should be evaluated in combination with water limitation (Ruehr et al., 2016). Under both, heat and drought-heat, stem diameter has been shown to level off or decrease, with larger effects occurring under water limitation (Bauweraerts et al., 2014; Ruehr et al., 2016). Large proportions (>90%) of stem diameter fluctuations result from swelling or shrinkage of the elastic bark, indicating tree water deficits (Zweifel et al., 2000; Zweifel and Häsler, 2000). The tree water status in combination with gas exchange measurements could reveal important evidence on tree physiological mechanisms and processes.

In addition to evaluating stress effects, information about recovery dynamics are crucial for estimating the resilience of trees and forests to climatic change in order to adapt long-term forest management. Stress impacts may leave physiological functions impaired for weeks and even years post-stress (Ruehr et al., 2019; Rehschuh et al., 2020; Schuldt et al., 2020). Depending on stress severity (Ruehr et al., 2019), this includes the persistent reduction of gas exchange (Duarte et al., 2016; Ruehr et al., 2016; Birami et al., 2018; Rehschuh et al., 2020), often associated with the impairment of the photosynthetic apparatus (Ameye et al., 2012; Birami et al., 2018) or the lack/only partial recovery of the hydraulic system (Brodribb et al., 2010; Rehschuh et al., 2020). Further, stem increment rates have been shown to remain low (Anderegg et al., 2015; Rehschuh et al., 2017), particularly due to the reduction of leaf area and C reserves during stress (Galiano et al., 2011). Therefore, the rapid recovery of net C gain plays an important role during recovery, as it provides energy and C skeletons for repair and regrowth processes.

Here we tested the impacts of a heatwave on the hydraulic and metabolic recovery of well-watered and drought-treated *Pinus sylvestris* seedlings. For this, seedlings were placed in individual gas exchange chambers and exposed to a heat and a hot drought scenario, followed by a recovery period, while continuously measuring above- and belowground CO_2_ and H_2_O gas exchange, and stem diameter change. Our objectives were to quantify the dynamics of (1) water relations during heat stress, in particular the cooling capacity of transpiration, (2) leaf temperature stress impact and recovery of leaf and branch hydraulic parameters, and (3) the interplay of hydraulic processes with carbon relations, and the effect on stem diameter change. We further investigated parameters related to functional integrity of leaves such as the maximum light-adapted quantum yield of the photosystem II (F′_v_/F′_m_) and electrolyte leakage. To provide a complete picture of carbon and water relations, we analyzed the diurnal dynamics within several temperature regimes, as well as the initial and final recovery phase of the experiment. The following hypotheses were tested:

1. Low transpiration rates during heat waves result in higher thermal stress leading to tissue damage.
2. Metabolic and hydraulic recovery will be fast if stress does not result in functional impairment and/or damage.
3. Stress-induced reductions of hydraulic properties will be directly linked to the recovery of the tree C balance and stem growth.

## Material and Methods

### Plant material and growth conditions

3-year-old *Pinus sylvestris* seedlings originating from a tree nursery in Middle Franconia (provenance 85115), Germany, were potted in round pots (18 cm in height, 22 cm in diameter) in March 2018. To assess the whole-tree C balance (net C uptake = net photosynthesis-respiration) we used a C-free potting substrate, i.e. a mixture of fine quartz sand (0.1-1.2 mm), medium-grained sand (1-2.5 mm), gravel (3-5.6 mm) and vermiculite in a relation of 2:2:1:2. The substrate was enriched with 12 g of slow-release fertilizer (Osmocote® Exact Standard 5-6M fertilizer 15-9-12+2MgO+TE, ICL Specialty Fertilizers, The Netherlands) per pot and liquid fertilizer (Compo® Complete, 6+4+6(+2) NPK(MgO), Hornbach, Germany) added monthly during the growing season. During the entire adjustment and experimental period, seedlings were kept in a scientific greenhouse facility in Garmisch-Partenkirchen, Germany (708 m asl, 47°28′32.9″N, 11°3′44.2″E) with highly UV-transmissive glass. Additionally, growth lamps (T-agro 400W, Philips, Hamburg, Germany) were used to supplement outside light. Photosynthetic active radiation (PAR) inside the greenhouse was measured continuously (PQS 1, Kipp & Zonen, Delft, The Netherlands), reaching daytime averages of 350 to 550 μmol m^−2^ s^−1^ (16 h daylight length).

Seedlings were randomly assigned to a well-watered control, a well-watered heat and a combined drought-heat treatment, in which we assessed stress and recovery trajectories. After needle elongation was completed (August 02, 2018), seedlings assigned to the drought-heat treatment were subjected to a 1.5-month drought period, keeping the soil water content (SWC) close to 10% (EC5, Meter Group, USA) before heat stress was initiated. SWC was adjusted automatically by a drip irrigation system (Rain Bird, Azusa, CA, USA), and was close to field capacity at 22% in the control treatment. Throughout the pre-drought period, air temperature and relative humidity (RH; CS215, Campbell Scientific, Logan, UT) were maintained constant (CC600, RAM Regel- und Messtechnische Apparate GmbH, Herrsching, Germany) between well-watered and drought treated seedlings (daytime: temp.: 22.95±0.07 °C; RH: 71.6±0.5%; nighttime: temp.: 17.36±0.02°C; RH: 81.8±0.8%).

On September 18, 2018, we started the experiment in the gas exchange chambers. Seedlings used for the control, heat and drought-heat treatment were on average 57.3±1.8 cm (±SE), 57.5±1.9 cm and 58.1±3.6 cm in height, and had a stem diameter of 16.4±0.6 cm, 15.9±0.5 cm and 14.9±0.4 cm, respectively.

To realistically adjust the environmental conditions in the stress treatments to extreme heat events, we analyzed daily and hourly data (air temperature, RH, VPD) from the *German Meteorological Service* of climatically extreme years in Weißenburg-Emetzheim, Franconia, Germany (435 m a.s.l., N49° 1′ 38,28″ E10° 58′ 55,2″). Around this area, Scots pine dieback had been observed after the hot dry summer in 2015 (Gößwein et al., 2017; Walentowski et al., 2017). As we found the period from August 07-13, 2003 to reveal the most intense climate extremes, we analyzed diurnal hourly averages for this period (Fig. S1) to base the climatic conditions in the tree chambers on these data. We stepwise increased the air temperature in the stress treatments over 20 days (30°C, 33°C, 35°C, 38°C and 40/41°C period) after acclimating the seedlings in the tree chambers for 5 d at 26°C, with VPD increasing concurrently to max. 6 kPa in the drought-heat treatment, but remaining lower (max. c. 4 kPa) in the heat treatment due to larger transpiration rates and thus higher humidity (Fig. 1a-b, Fig. 2a). Air temperatures were maintained at the targeted maximum values for min. 6 h per day. Soil temperature increased with a timely offset to air temperature, and also increased slightly in the control treatment, because soil compartments could only be cooled passively (Fig. 1c). During night time, air and soil temperatures decreased to c. 15-20°C in all treatments.

**Fig. 1:**
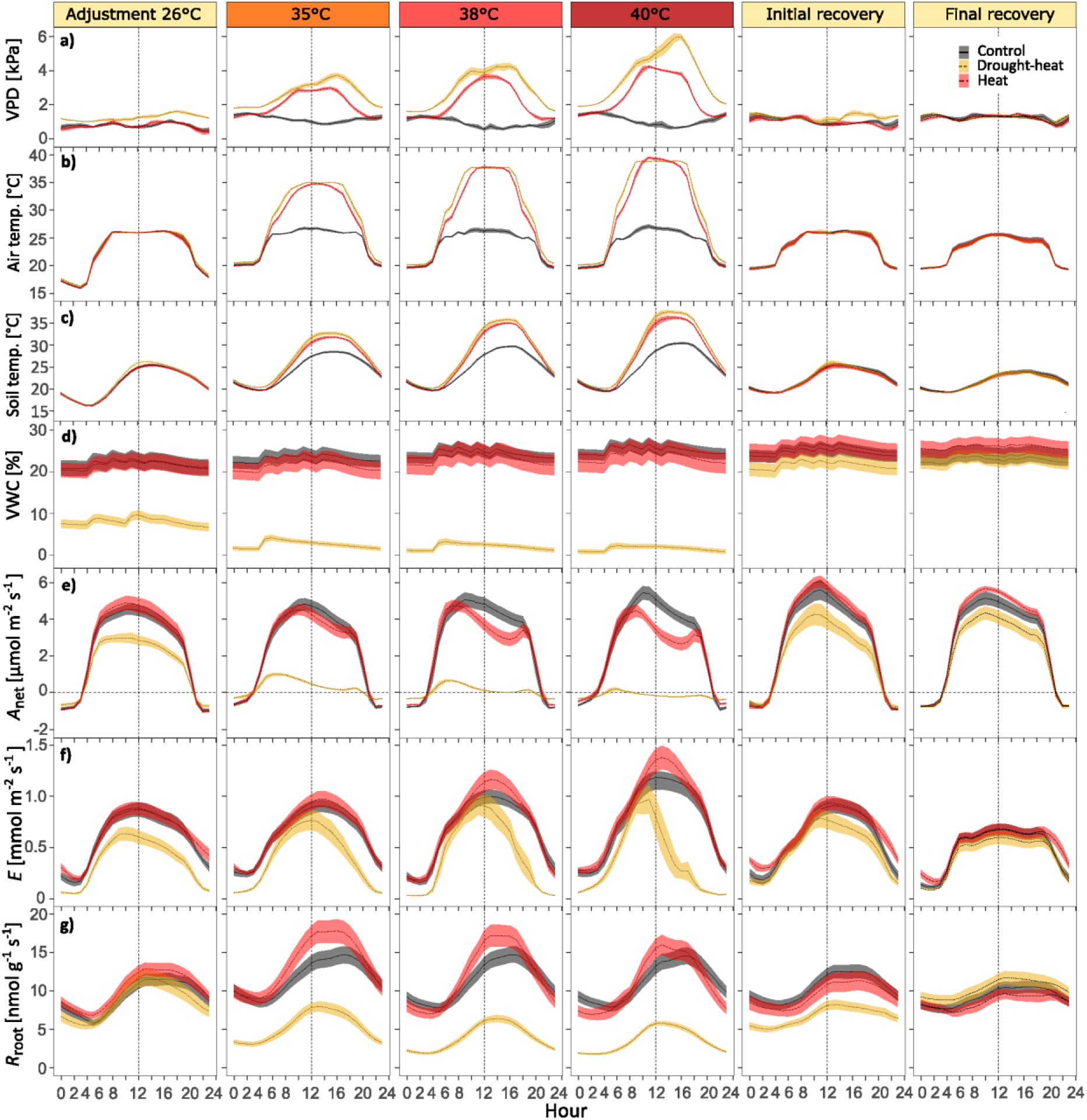
Diurnal dynamics of treatment-averaged hourly a) vapor pressure deficit (VPD), b) air temperature c) soil temperature, d) soil volumetric water content (VWC), e) net assimilation (*A*_net_), f) transpiration (*E*), and g) root respiration (*R*_root_) for different periods. Before heat stress, during adjustment (26°C, 4 days), temperature increments (35°C, 7 days; 38°C, 3 days and 40°C, 4 days), the initial (26°C, day 28-30) and final recovery period (26°C, day 40-42). Daytime length was 16 h (4:30 – 20:30 CET). Shown are treatment averages and shaded areas are ±SE (n = 6 per treatment).

**Fig. 2:**
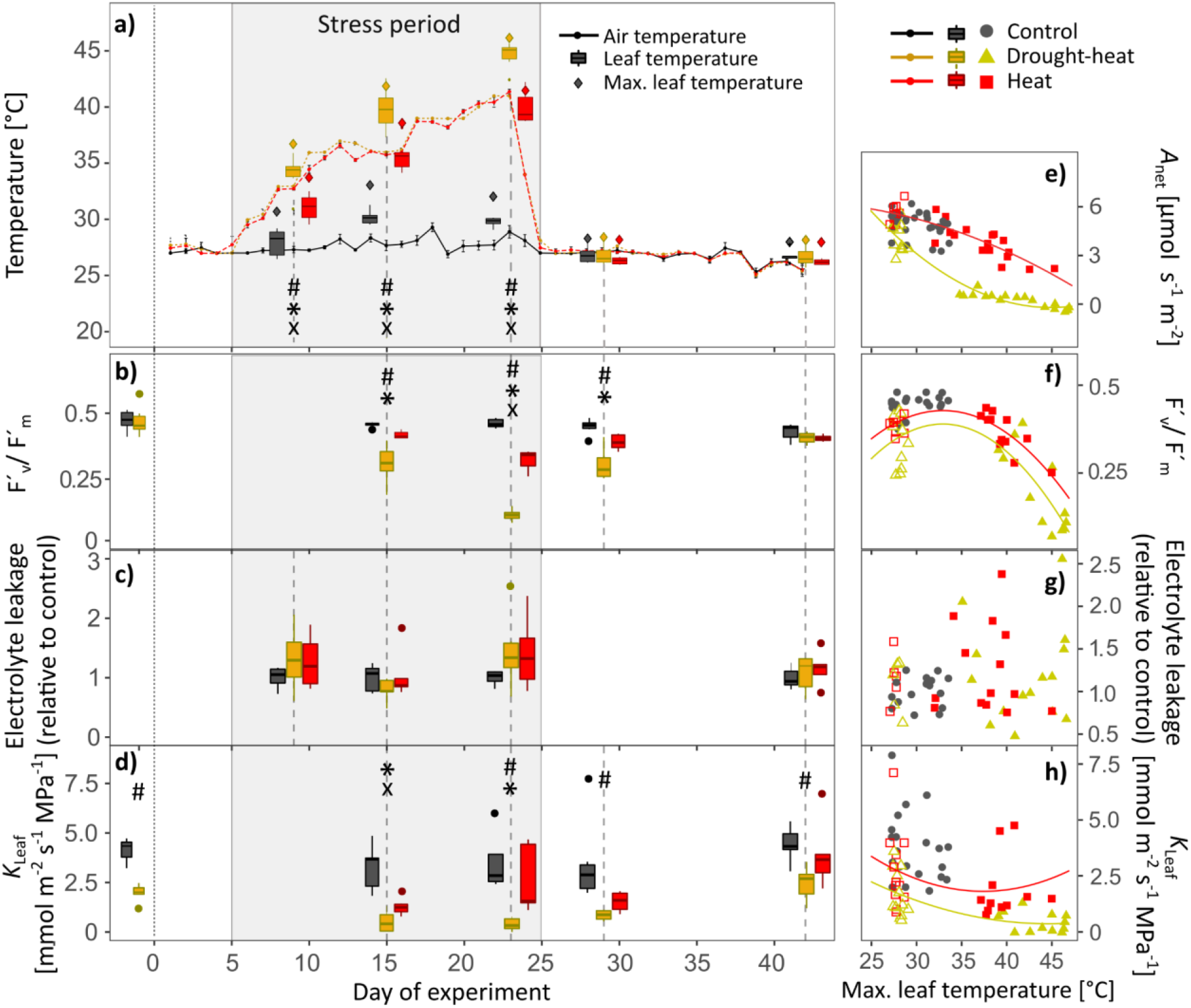
Dynamics of air and leaf temperature and stress indicators during the experimental phase in Scots pine seedlings. Shown are a) air temperature, mean and maximum leaf temperature, b) maximum light adapted quantum yield of photosystem II (F′_v_/F′_m_), c) electrolyte leakage and d) leaf hydraulic conductance (K_Leaf_) during stress progression and recovery (n = 5-6 per treatment). In e-h) the relationships of A_net_, F′_v_/F′_m_, electrolyte leakage and K_Leaf_ with maximum leaf temperature during stress and recovery are shown per seedling and treatment. The gray boxes represent the stress period. Symbols indicate significant differences (P<0.05) between treatments per sampling campaign (Kruskal-Wallis and Bonferroni post-hoc test) as follows: drought-heat vs. control (#), heat vs. control (x), and drought-heat vs. heat (⚹). Note that data before day 0 are for measurements 2 weeks before the chamber experiment started and that drought-heat seedlings were drought pre-stressed for 1.5 months. Open symbols in e-h) indicate measurements during the recovery period. Polynomial curves were fitted for the drought-heat and heat treatment if applicable (for functions see Supplemental Table S1).

The conditions in the control treatment were adjusted according to August 30-year-averages (1986-2016) in Weißenburg-Emetzheim, and maintaining SWC close to field capacity by watering 4x daily c. 75 ml, respectively (Fig. 1d). A similar SWC was achieved in the heat treatment, while the amount had to be increased slightly due to higher transpiration rates. In the drought-heat treatment, seedlings were irrigated 2x daily during acclimation (c. 50 ml, respectively) and only at 5 am during the heat stress period (70 ml), while the water supply was further steadily reduced.

To analyze recovery processes, seedlings were re-watered to field capacity at 5 am on October 13, 2018, and temperature and VPD were down-regulated to control conditions (Fig. 1a-d, Fig. 2a, see initial and final recovery period).

### Tree gas exchange chambers

For continuous monitoring of above- and belowground gas exchange (H_2_O and CO_2_) during the drought and heat period and the recovery phase, we randomly selected 12 well-watered (subsequently assigned to n=6 control and n=6 heat treated) and 6 drought pre-treated seedlings (assigned to the drought-heat treatment), and placed them in individual tree chambers within the greenhouse compartment. The tree chambers were divided into a translucent temperature-controlled shoot compartment, which was gas-tightly sealed from an opaque root compartment (for further details on materials and technical operation of the chamber system see Birami et al. 2020). Air temperature was predefined for the different treatments and days and regulated within each shoot compartment separately by fast-response thermocouples (5SC-TTTI-36-2M, Newport Electronics GmbH, Deckenpfronn, Germany) and based on a water-cooling principle. PAR was recorded per chamber by photodiodes (G1118, Hamamatsu Photonics, Hamamatsu, Japan), calibrated in advance with a PAR sensor (PQS 1, Kipp & Zonen, Delft, the Netherlands). Soil temperature and moisture were continuously logged (TS 107, Campbell Scientific, Inc. USA and EC 5, Meter Group, USA, respectively), and data recorded in 10 min intervals (CR1000, Campbell Scientific, Inc. USA). In addition, stem diameter change was measured half-hourly by dendrometers (DD-S, Ecomatik, Dachau, Germany), attached to each stem at approximately 5-10 cm above the soil surface.

### Gas exchange measurements and calculations

Above- and belowground chamber compartments were constantly supplied with air (Air_supply_) from an oil-free screw compressor by adding a pre-defined CO_2_ (c. 430 ppm) and H_2_O concentration (8 mmol during the adjustment and recovery period; 4 mmol during the stress period). Each of the 20 shoot and root compartments received c. 13 L min^−1^ and 3 L min^−1^ of air flow, respectively.

Firstly, absolute [CO_2_] and [H_2_O] of Air_supply_ and sample air (Air_sample_) were quantified by a gas analyzer (Li-840, LI-COR, Lincoln, NE, USA). Subsequently, a differential gas analyzer determined the differences between Air_supply_ and Air_sample_ (Li-7000, LI-COR, Lincoln, USA). We used two empty tree chambers (no trees but pots with the same potting substrate) to correct for fluctuations in [CO_2_] and [H_2_O] not caused by plant gas exchange. Subsequently, offsets were removed from the data. Every compartment was measured once every 2 h for c. 80 sec, while data was logged every 10 sec. The following equations were used to calculate gas exchange fluxes: Transpirational loss of H_2_O (*E*) in mol m^−2^ s^−1^ was calculated as follows:

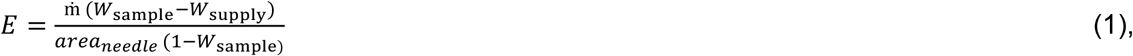

where ṁ is air mass flow (mol s^−1^), *W*_supply_ is [H_2_O] in Air_supply_, and *W*_sample_ is [H_2_O] in Air_sample_. Area_needle_ is the total needle area of the shoot in m^2^.

Stomatal conductance *g_s_* in mmol m^−2^ s^−1^ was calculated accordingly:

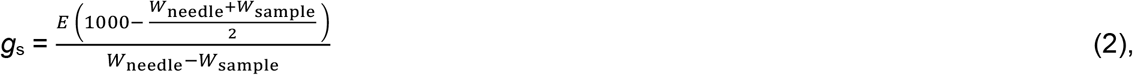

where *W*_needle_ is the needle H_2_O vapor concentration, which was deduced from saturated vapor pressure (kPa) at the prevailing air temperature (°C) and atmospheric pressure p (kPa). This method of determining *g*_s_ assumes well mixed air within the shoot compartment and therefore, neglects the boundary layer conductance.

Net assimilation (*A*_net_) and soot respiration (*R*_shoot_) in μmol m^−2^ s^−1^ and root respiration (*R*_root_) in μmol s^−1^ g ^−1^ DW (dry weight) were calculated by the equation for CO_2_ fluxes:

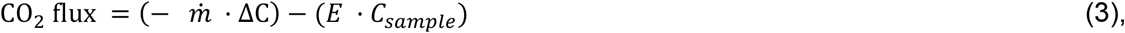

where ΔC is the difference between [CO_2_] of Air_supply_ and Air_sample_, *C*_sample_ is [CO_2_] of Air_sample_. *E* was included to correct for dilution by transpiration. Further, *A*_net_ was related to total needle area, and *R*_root_ to *g* DW of total root biomass. Whole-tree daily net C uptake (in mg C d^−1^ tree^−1^) was then calculated from daily sums (calculated by summing up hourly means) by subtracting daily C loss (*R*_root_ and *R*_shoot_) from daily C uptake via assimilation:

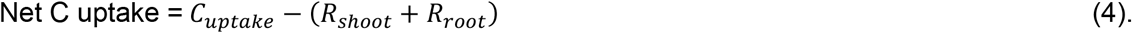

### Leaf temperature

To assess temperature and indirect drought effects, we measured leaf temperature (n=6 per treatment) between 1 and 2 pm using an infrared camera (PI 450, Optris, Germany). Leaf temperature was evaluated about weekly during the stress period, and two times during the recovery phase (initial and final recovery). Mean and maximum leaf temperatures were determined from images using the manufacturer’s software. For this, we corrected for background radiation using air temperature during measurements and set emissivity to 0.97.

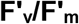

We measured maximum light adapted quantum yield of photosystem II (F’_v_/F’_m_) using a portable leaf-gas exchange system (Li-6400, LI-COR Inc., Lincoln, NE, USA), supplemented with a fluorescence head (6400-40 Leaf Chamber Fluorometer). Measurements (n=6 per treatment) were conducted between 9 am and 1 pm about 2 weeks before placing the seedlings in the tree chambers (only drought stress), during drought and/or heat stress and during the recovery phase, coordinated with leaf temperature measurements. For this, we clamped needles into the leaf cuvette, covering it completely (2 cm^2^), but avoiding overlapping of needles as much as possible. We used pre-determined saturated light conditions of 1200 μmol m^−2^ s^−1^ PPFD and a reference [CO_2_] of 400 ppm. Leaf temperature, RH and VPD were kept constant during measurements, and averaged 26°C, 52% and 1.5 kPa in the control and 29°C, 37% and 3 kPa in the heat and drought-heat treatment.

### Electrolyte Leakage

To assess cell membrane stability, we quantified electrolyte leakage of needles following Mena-Petite et al. (2003). Sampling of 10 needle fascicles on 5 seedlings per treatment was done in coordination with leaf temperature measurements, between 10 and 12 am. The samples were immediately wrapped in moist paper towels and subsequently washed in a petri dish filled with deionized water to remove solutes from needle surfaces. Excess water was removed and needles were cut into segments of c. 1 cm length, excluding the needle sheath. The needle segments were weighed and transferred to falcon tubes (Isolab, Laborgeräte GmbH, Eschau, Germany) with 20 ml of distilled water of known electrical conductivity (EC_dist_). Electrical conductivity was determined with a conductivity electrode (1480–90 Cole-Palmer Instruments; Chicago, I., USA). Subsequently, falcon tubes were placed inside a desiccator. To facilitate electrolyte leakage, a vacuum was created by withdrawing air. Samples were kept in the dark for 24 h at room temperature. Then, samples were shaken and EC_final_ was measured. Subsequently, samples were heated to 100°C for 10 min to destroy all living cells and maximize leakage. After samples were cooled to room temperature, EC_total_ was determined. 24 h- electrolyte leakage was then calculated as follows:

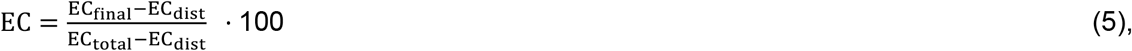

which was then related to needle weight and expressed relative to control seedlings.

### Leaf hydraulic conductance (*K*_Leaf_)

*K*_Leaf_ was measured between 10 and 12 am on 5-6 seedlings per treatment, coordinated with F’_v_/F’_m_ measurements. We adjusted the evaporative flux method used by Sack and Scoffoni (2012) to measure single needles of seedlings as follows. Needle fascicles were detached from branches under water and placed with their base in Eppendorf tubes filled with distilled and filtered water (0.22 μm pore size). After an adjustment time of c. 20 min in ventilated air to enhance *E* under constant light conditions (800-1000 μmol m^−2^ s^−1^ PAR), *E* of a single needle fascicle was measured using a portable leaf-gas exchange system (Li-6400, LI-COR Inc., Lincoln, NE, USA) equipped with a light source (6400-40 Leaf Chamber Fluorometer) under saturated light conditions of 1200 μmol m^−2^ s^−1^ PAR and [CO_2_] of 400 ppm until stable conditions were reached. Temperature and RH during acclimation and measurement corresponded to conditions inside the tree chambers per treatment and measurement time point. After measuring *E*, the same needle fascicles were placed in nontransparent plastic bags to equilibrate for 20 min whereupon *ψ*_Needle_ was determined using a Scholander pressure chamber (Model 1000, PMS Instruments, Albany, Oregon, USA). The projected needle area (*A*_Leaf_) of the enclosed needle segments was determined using a leaf area meter (Li-3100, LI-COR Inc., Lincoln, Nebraska, USA). *K*_Leaf_ in mmol m^−2^ s^−1^ MPa^−1^ was then calculated as follows:

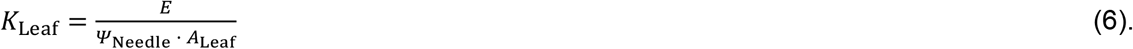

### Needle water potential (*ψ*_Needle_) and relative needle water content (RWC_Needle_)

To assess tree water status, we measured predawn and midday *ψ*_Needle_ of mature needles of 6 seedlings per treatment using a Scholander pressure chamber (Model 1000, PMS Instruments, Albany, Oregon, USA). We report daily minimum *ψ*_Needle_ since predawn *ψ*_Needle_ was measured before the automatic irrigation and hence sometimes slightly lower than midday *ψ*_Needle_. Measurements of *ψ*_Needle_ were coordinated with RWC_Needle_ measurements. For this, we sampled two needle fascicles per seedling (n=6 per treatment), placed them in a plastic bag to avoid evaporation and immediately determined fresh weight (*W*_fresh_). After soaking them in purified water at room temperature for 48 h (as predetermined from saturation curves), turgid weight (*W*_turgid_) was measured, then needles placed in an oven at 70°C for 48 h, and consequently dry weight (*W*_dw_) determined. RWC_Needle_ was calculated as follows:

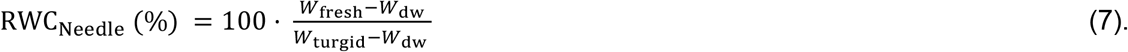

### Biomass and needle area

We sampled the biomass of the seedlings (n=6 per treatment) at the end of the experiment and took a subsample to determine stem hydraulic conductivity (see below). Biomass was then separated into leaf, root, branch and stem tissues and oven-dried for 48 h at 70°C to determine dry weight.

Leaf area of each tree was derived from leaf biomass and pre-determined specific leaf area (gDW cm^−2^). On a subsample of fresh needles, specific leaf area was measured using an area meter (Li-3100, LI-COR Inc., Lincoln, Nebraska, USA), and its dry mass determined. Total leaf area per tree was then calculated via this leaf area index.

### Stem xylem hydraulic conductivity (*K*_S_)

A part of each stem was wrapped in cling film and frozen immediately in plastic bags at −18°C to determine *K*_S_ (n=6 per treatment) by applying the pressure-flow method (Sperry and Tyree, 1988), as described more detailed in Rehschuh et al. (2020). In short, frozen samples were thawed in distilled water, cut back underwater with sharp instruments, and the bark was peeled off. Samples were fixed in a fivefold valve (Luer-lock system; neoLab Migge Laborbedarf-Vertriebs GmbH, Heidelberg, Germany), and exposed to a natural pressure of water flow to determine in-situ degree of embolism. A mass flow meter (mini-CORI-FLOW M13, Bronkhorst, Montigny les Cormeilles, France) served to measure the flow rate of each stem segment individually. By considering the xylem cross-sectional area and sample length, we then determined *K*_S_.

### Statistical data analysis

Gas exchange data were quality-checked per chamber, and outliers identified separately for day- and nighttime measurements based on the widely applied boxplot approach. For this, values outside 1.5 times the interquartile range above the upper and below the lower quartile were removed. Between day 32 and 34, daily net C uptake was calculated from linear interpolation due to data unavailability because of technical constraints. For diurnal progression analysis of gas exchange data and depiction of environmental variables, we hourly averaged data from several key experimental periods (adjustment period at 26°C, temperature stages (35°C, 38°C and 40°C), initial stable and final recovery period).

Data processing and statistical analyses were performed in ‘R’ version 3.6.1 (R Core Team, 2019). Differences between treatments (significant if P<0.05) were determined per measurement campaign for discretely measured parameters (e.g. leaf temperature, F′_v_/F′_m_, *K*_Leaf_) by applying the Kruskal-Wallis followed by the Bonferroni post-hoc test, therefore accounting for small sample sizes.

Treatment effects on continuous data (gas exchange measurements, net C uptake) were assessed by applying linear-mixed effects models (lme; lmerTest package; Kuznetsova et al., 2017). Treatment and time period were assigned as fixed effect and chamber as random factor. Time periods included the adjustment period, the different temperature steps, initial and final recovery. For diurnal measurements, we further considered day- and nighttime. We selected the model with the lowest Akaike’s information criterion corrected for small sample size (AICc; Burnham and Anderson, 2002), i.e. the most parsimonious model. Further, we used the post-hoc Tukey’s multiple comparisons test of means for the determination of differences between treatments (package emmeans; Lenth et al., 2020). Tukey′s HSD is reported here.

Further, we analyzed leaf temperature dependencies of *A*_net_, F′_v_/F′_m_, electrolyte leakage and *K*_Leaf_, and relationships of *ψ*_Needle_ with gas exchange parameters (*A*_net_, net C uptake, *E*). Additionally, we tested for the relationships of net C uptake with stem diameter change during stress and recovery separately. Where applicable, we fitted polynomial or sigmoidal functions for better visualization of relationships. Fitted curves for stem diameter change in relation to C uptake during recovery were analyzed for significant differences between treatments by comparing their coefficients.

## Results

Drought stress was initiated after needle growth was completed and no significant differences between treatments in needle biomass and area were found (Fig. S2). However, a modest effect of the prolonged drought became visible as the root:shoot ratio (−25%), root (−23%) and woody biomass (−15%) tended to be lower than in control seedlings.

### Tree gas exchange, net C uptake and stem diameter change during stress and recovery

During daytime, increases in air temperature enhanced VPD (Fig. 1a-b). This increase in VPD was larger in the drought-heat treatment due to lower transpiration rates, which reduced the RH inside the tree chambers. Nighttime temperatures, however, did not differ largely between treatments, and were maintained close to 20°C.

The diurnal course of *E* and *A*_net_ in drought-treated seedlings remained below the control already before the heatwave was initiated (Tukey’s HSD, P<0.01, see Table S2 for p-values; Fig. 1e-f). With increasing temperatures and VPD, *E* in drought-heat seedlings approximated control values during the morning hours, but decreased dramatically in the course of the day, intensified with stress progression (Tukey’s HSD, P<0.05 for 38°C and P<0.001 for 40°C period). In well-watered heat treated seedlings, *E* increased with a rise in air temperature and VPD, despite declining *g*_s_ (Fig. S3b). This was reflected in diurnal dynamics of *A*_net_, which decreased to about 70% of control values during periods of high air temperature and VPD (Tukey’s HSD, P<0.001 for 40°C period; Fig. 1a-b) between 10 am and 4 pm, followed by a slight increase later (Fig. 1e). Under the combined drought and heat stress, we found a strong decline in *A*_net_, which was reflected in partial or full stomatal closure (Fig. S4), with *g*_s_ being lower at the same VPD and air temperature compared to the heat treatment (Fig. S3).

*R*_root_ largely followed diurnal soil temperature patterns (Fig. 1c,g, Fig. S5). Over the 20-day heat period, *R*_root_ initially increased under well-watered conditions, but when 40°C prevailed, the difference to control seedlings diminished, indicating some kind of adjustment. In the drought-heat treatment, *R*_root_ declined with an increase in soil temperatures and was about three times lower than in the control during day- and nighttime throughout the 35°C to 40°C period (Tukey’s HSD, P<0.01; Fig. 1g).

The daily net C uptake appeared to be moderately sensitive to heat under well-watered conditions, particularly when air temperatures reached >38°C and VPD rose >3 kPa (Fig. 3a, see also Fig. S6 for daily means of *A*_net_, *R*_shoot_ and *R*_root_ per tree). The effect on the C balance was more apparent when heat was combined with drought. While net C uptake and stem growth (Fig. 3a-b) continued under heat, net C uptake decreased in drought-heat treated seedlings, resulting in a negative C balance (Tukey’s HSD, P<0.001 compared to heat and control). This was reflected in stem growth cessation and phloem shrinkage due to water deficits (Fig. 3b).

**Fig. 3:**
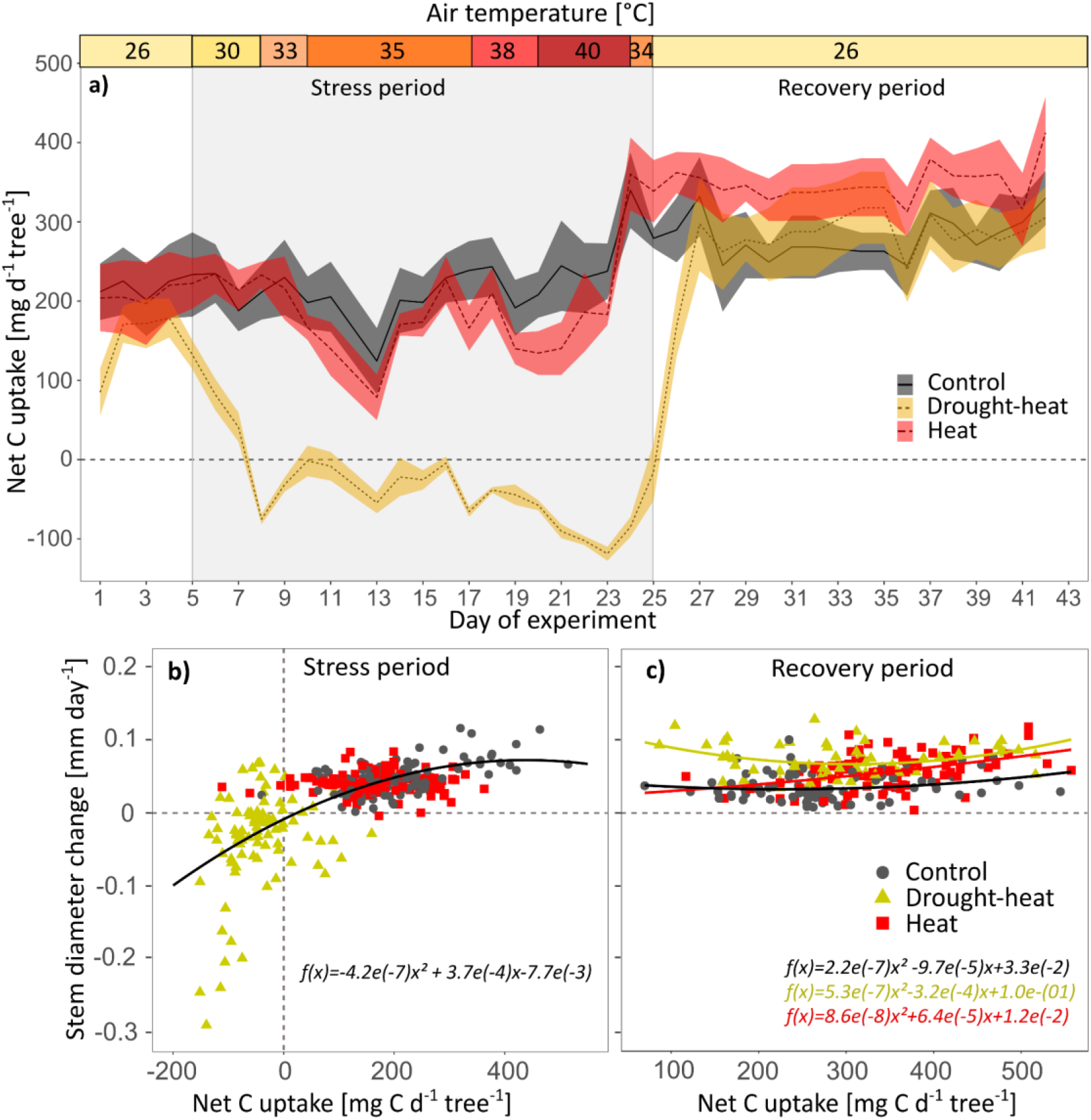
Daily net carbon (C) uptake of Scots pine seedlings during the entire experiment (a) and in relation to stem diameter change during stress (b) and recovery (c). The daily net C uptake per seedling was derived by summing hourly means of C uptake by photosynthesis minus C release by shoot and root respiration. In a), the shaded areas show ±SE (n = 6). Mean daytime air temperature is given for the stress treatments, while control seedlings were at 26-28°C. In b) and c) diurnal stem diameter changes per seedling and treatment are shown. Negative stem diameter change was caused by stem shrinking due to water deficits. The rehydration period after stress release (6 days) was excluded for the recovery period.

With stress release on day 25 of continuous gas exchange measurements, we observed a fast response of *E* and net C uptake (Tukey’s HSD, P>0.1 for stress treatments compared to control; Table S2). In the heat treatment, net C uptake tended to exceed that of the control seedlings by c. 10%, also reflected in higher daily stem increment rates (P>0.05; Fig. 3c). Net C uptake in drought-heat treated seedlings reached control values within 2 days, while *A*_net_ remained c. 20% below control rates (Tukey’s HSD, P<0.05 for initial and P=0.08 for final recovery period; Fig. 1e). The fast recovery of tree net C uptake can be explained by an initially slow recovery of *R*_root_ (Tukey’s HSD, P<0.05 compared to control, Fig. 1g) and a tendency for a lower root:shoot ratio (i.e. *R*_root_ at the tree level did not fully recover, see Fig. S6). Alongside, stem growth rates exceeded those of the control during the second half of the recovery period largely independent of net C uptake (P<0.05; Fig. 3c).

### Stress impairment and delayed recovery

Because the degree of evaporative cooling from transpiration differed among treatments, seedlings exposed to combined heat and drought experienced substantially (3-4.5°C) higher leaf temperatures than heat treated well-watered seedlings at the same air temperature (Fig. 2a, Fig. S7). Leaf temperatures were significantly higher compared to the control in both stress treatments, while in well-watered heat treated seedlings evaporative cooling maintained leave temperature below maximum air temperature.

The effect of high leaf temperatures was also apparent in the decline of *A*_net_ and simultaneous decrease in photosystem II activity (here F′_v_/F′_m_; Fig. 2e-f). F′_v_/F′_m_ was not affected by the pre-drought, but decreased strongly in response to heat and water limitation within the course of the experiment, reaching 80% lower values than the control (P<0.05; Fig. 2b). Heat treated seedlings were less affected, but F′_v_/F′_m_ also decreased by 30% compared to the control (P<0.05). These effects were fully reversed within the recovery period in both stress treatments, with a delay in drought-heat seedlings. While the relationship between *A*_net_ and leaf temperature differed under heat and drought-heat (including recovery), F′_v_/F′_m_ showed a rather similar response pattern to leaf temperature within the two stress treatments (Fig. 2f). In contrast to large stress effects on photosystem II, we found no apparent needle cell damage from heat stress on well-watered and drought-treated seedlings as electrolyte leakage only tended to be increased on day 23 when highest leaf temperatures were reached, but did not differ significantly from the control (Fig. 2c,g).

Further, hydraulic parameters were affected mildly under well-watered conditions, but strongly under water limitation. *K*_Leaf_ was reduced during the initial drought period, and continued to decline with heat initiation in both stress treatments (Fig. 2d), reaching c. 90% lower values in the drought-heat treatment compared to the control (P<0.05). This decline was partially associated with high leaf temperatures (Fig. 2h). While the other stress-related parameters recovered fully, *K*_Leaf_ did not reach control values in drought-heat treated seedlings until the end of the experiment (P<0.05). Similarly, *K*_S_ was c. 25% lower post drought-heat compared to the control at the end of the experiment, indicating that *K*_S_ did not recover (P<0.05, Fig. 4a-b). However, the stress impacts were less apparent than for *K*_Leaf_ (Fig. 4c).

**Fig. 4:**
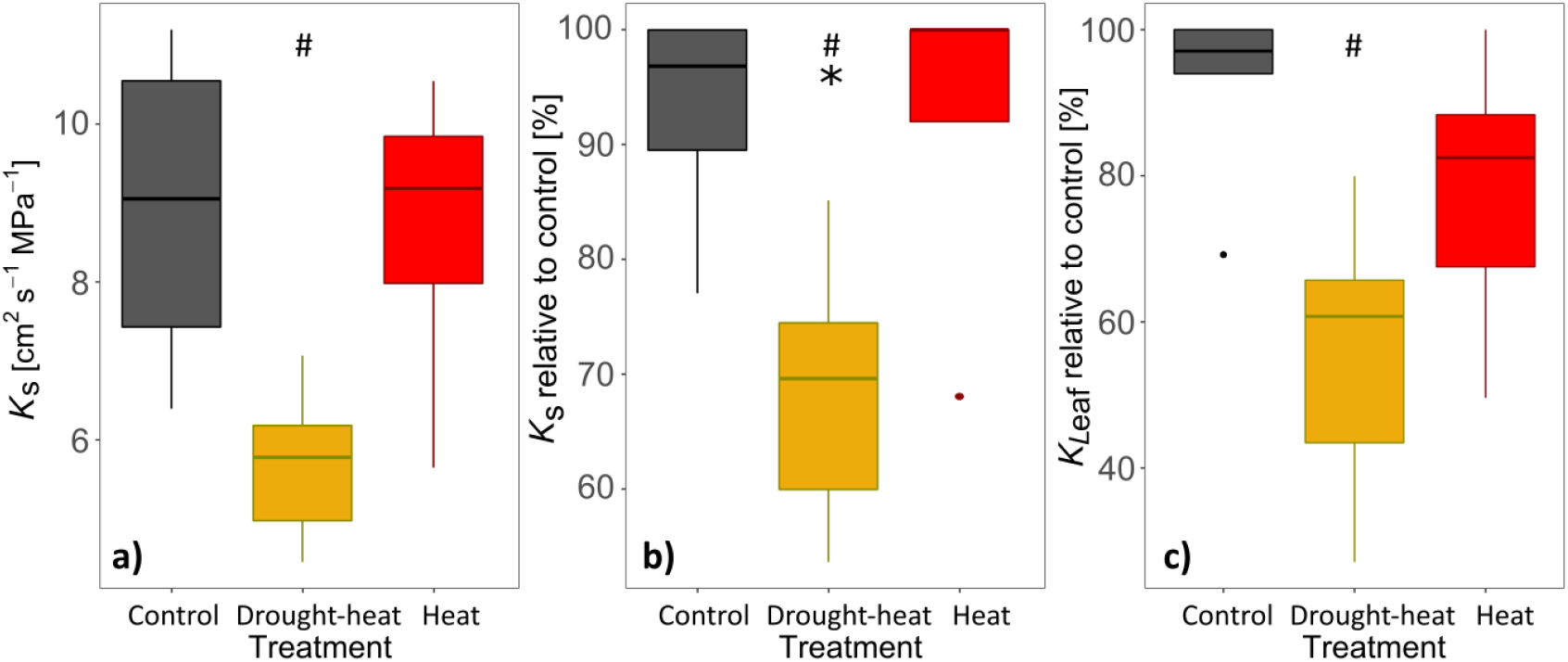
a) Treatment-specific stem hydraulic conductivity (K_S_), b) K_S_ relative to average control values, and c) leaf hydraulic conductance (K_Leaf_) relative to average control values measured on the last day of the recovery period, 18 days after re-watering (n=5-6). Symbols indicate significant differences (P<0.05) between drought-heat and control (#), and between drought-heat and heat (⚹) (Kruskal-Wallis & Bonferroni post-hoc test).

The combination of drought and heat stress had large impacts on *ψ*_Needle_, which decreased to −2.70±0.13 MPa (P<0.05, Fig. 5a). While heat stress alone did not affect *ψ*_Needle_, RWC_Needle_ declined significantly in the heat treatment compared to the control during the hottest days (Fig 5b). In the drought-heat treatment, the decline in RWC_Needle_ was more pronounced and appeared earlier during the experiment. *ψ*_Needle_ and RWC_Needle_ recovered to control values nearly immediately following stress release.

**Fig. 5:**
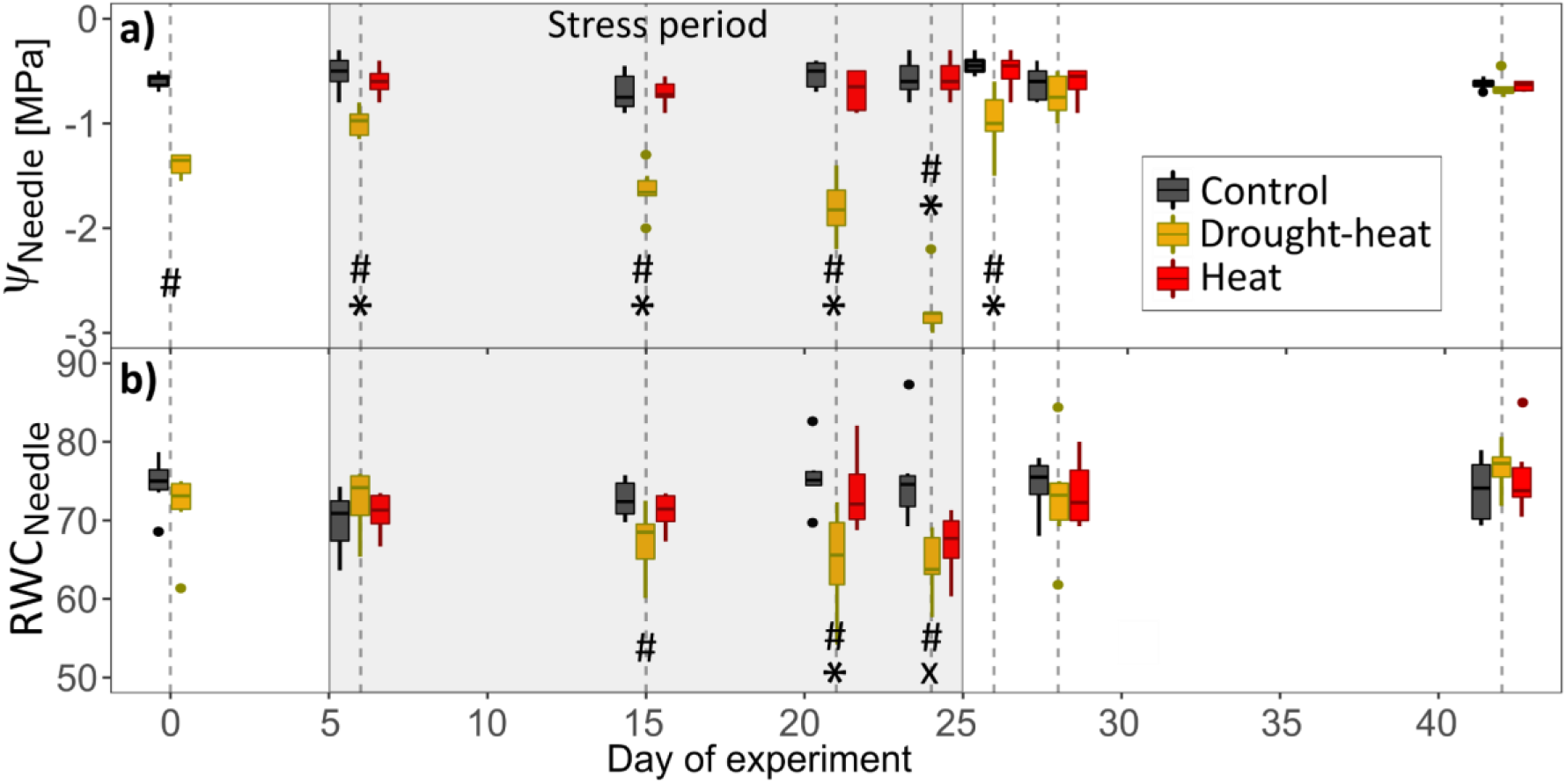
Treatment-specific dynamics of a) needle water potential (ψ_Needle_, n=6) and b) relative needle water content (RWC_Needle_, n=6) during stress and recovery. The gray boxes represent the stress period. Symbols indicate significant differences (P<0.05) between treatments per measurement campaign (Kruskal-Wallis and Bonferroni post-hoc test) as follows: drought-heat vs. control (#), heat vs. control (x), and drought-heat vs. heat (⚹). Note that drought-heat seedlings were drought pre-stressed for 1.5 months before the start of the main experiment (day 1).

Further, a close relationship of *A*_net_, net C uptake and *E* with *ψ*_Needle_ was apparent in the drought-heat treatment (Fig. 6). *A*_net_ halted or turned negative (i.e. shoot respiration and photorespiration exceeded assimilation) at a *ψ*_Needle_ of c. −1.16 MPa, while the tree net C uptake turned negative slightly earlier at a *ψ*_Needle_ of c. −1.08 MPa. This indicates the dominant role of tree hydraulic processes in limiting gas exchange rates during hot droughts. The recovery of these gas exchange parameters appeared to largely follow the increase in *ψ*_Needle_, with a tendency of somewhat lower *E* in previously drought-heat treated seedlings.

**Fig. 6:**
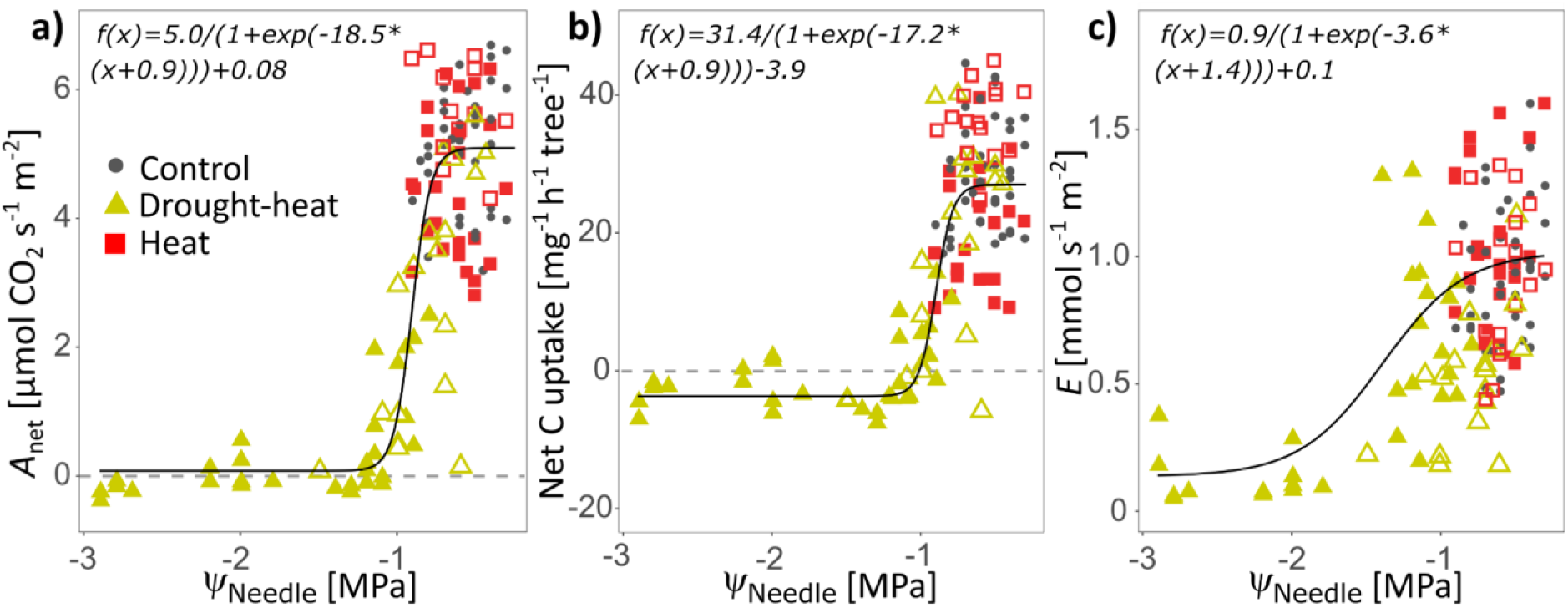
Dependencies of a) net assimilation (*A*_net_), b) net C uptake, and c) transpiration (*E*) with needle water potential (*ψ*_Needle_) measured at noon per seedling during stress and recovery. The different treatments are highlighted and open symbols indicate measurements during the recovery period. Sigmoidal functions were fitted to drought-heat (excluding recovery) and control measurements. Root-mean-square error:*A*_net_ 0.86; Net C uptake: 6.15;*E*: 0.30.

## Discussion

We closely monitored stress and recovery responses of Scots pine seedlings exposed to heat and hot drought stress. All seedlings successfully survived the heatwave, while additional drought delayed recovery as observed in photosynthetic and hydraulic responses. Moreover, we found all seedlings to recover overall net C uptake quickly, indicating that a new equilibrium between C uptake and release appeared largely independent of lasting hydraulic impairment.

### Thermal stress but no needle damage

A gradually increasing heatwave over 20 d resulted in daytime temperatures above 40°C on 3 d. Heat stress under well-watered conditions affected the hydraulic system moderately, but in combination with drought, stress impacts intensified. Both, on a diurnal scale and in the course of the heatwave, we found *E* to increase with rising air temperatures in well-watered seedlings (Fig. 1f; Ameye et al., 2012; Ruehr et al., 2016). Under water-limiting conditions and high VPD, *E* decreased strongly during the second half of the day to prevent extensive water loss and high xylem tensions, alongside declining *g*_s_ (Fig. S3; Ruehr et al., 2016; Birami et al., 2018; Zhao et al., 2013). This is in contrast to Urban et al. (2017b, 2017a), who found *g*_s_ to increase with rising temperatures, even under low soil water content. Stomatal adjustment affects evaporative cooling of transpiring leaves, thus decreasing leaf temperatures. In our study, well-watered heat seedlings were able to keep leaf temperatures below maximum air temperatures (Fig. 2a), as *E* increased despite declining *g*_s_. Under drought-heat, the limited capacity of needle cooling due to reduced *E* resulted in higher leaf temperatures (max. 46°C) during early afternoon. Although tree mortality has been reported following hot drought stress (Allen et al., 2010; Balducci et al., 2013; Teskey et al., 2015) and leaf temperatures similar to our study (47°C in Birami et al. (2018)), we did not find negative effects on seedling survival nor direct tissue damage, which was also supported by negligible effects on electrolyte leakage in needle tissues (Fig. 2c,g). In part, this might have been prevented by relatively high *E* rates during early mornings in drought-heat seedlings, buffering some of the excessive needle temperature stress. Moreover, the duration of the temperature exposure is important, while the heatwave resulted in 18 h at c. 40°C in our study. However, in well-watered seedlings of *Pinus radiata* at air temperatures and heat exposure comparable to our study, significant effects on the increase of electrolyte leakage were observed (Escandón et al., 2016). The discrepancy might result from the gradual increase of temperatures in our study compared to the immediate temperature increment from 25°C to 40°C in Escandón et al. (2016), allowing the seedlings to acclimate cell membrane stability to thermal stress (Saelim and Zwiazek, 2000).

While it is difficult to clearly disentangle the impacts of high temperatures on our drought treatment, we did not find direct needle damage albeit the very high leaf temperatures during periods of low transpiration (early afternoon). Hence, we found a remarkable thermal resistance of Scots pine needles to temperatures >45°C and have to partially reject our first hypothesis.

### Tree hydraulics during stress and recovery

Both, the heat and drought-heat treatment affected *K*_Leaf_ (Fig. 2d). In response to heat alone, the decline was modest and could be related to the high temperatures and/or increases in VPD. Most likely, heat stress affected *K*_Leaf_ in extra-xylary tissues due to tissue shrinkage, cell collapse and/or the reduction in aquaporin activity (Cochard et al., 2004; Scoffoni et al., 2014; Sack et al., 2016), consistent with the decline of RWC_Needle_ (Fig. 5b). In the drought-heat treatment, *K*_Leaf_ declined further alongside large reductions in *ψ*_Needle_ and RWC_Needle_, indicating that leaf xylem embolism must have appeared later during stress progression (Lo Gullo et al., 2005; Brodribb and Cochard, 2009; Johnson et al., 2009). The role of embolism was further supported by 25% reduced *K*_S_ at the end of the experiment in drought-heat seedlings (Fig. 4a-b). That stem xylem embolism reversal does not occur in Scots pine following drought release has been reported recently (Rehschuh et al., 2020). We also suggest embolism as underlying reason for the incomplete recovery of *K*_Leaf_ within the 18d-recovery-period (Fig. 4c). If solely extra-xylary tissues had been affected, we could assume *K*_Leaf_ to recover faster alongside RWC_Needle_ and *ψ*_Needle_ (Brodribb and Cochard, 2009; Laur and Hacke, 2014) due to hydration not related to refilling of xylem embolism. In agreement, in previously heat treated seedlings, we observed a fast recovery of RWC_Needle_ and *K*_Leaf_, supported by full *K*_S_ integrity (Fig. 4a-b). This agrees with a recent stress-recovery framework by Ruehr et al. (2019), indicating that xylem embolism delays hydraulic recovery, while outside xylem conductance can be recovered quickly. Additionally, the larger response of *K*_Leaf_ than *K*_S_ in response to water limitation indicates the higher vulnerability of distal organs such as leaves compared to stems (Brodribb and Cochard, 2009; Bartlett et al., 2016) in highly segmented trees such as Scots pine.

Overall, we show that under heat stress the impact on the hydraulic system became particularly apparent in drought-treated seedlings, resulting in needle and stem xylem embolism, which did not recover fully.

### Metabolic responses during stress and recovery

The stress imposed by high temperatures was further reflected in the C metabolism of the seedlings. *A_net_* declined moderately in response to higher leaf temperatures (c. 40°C) in well-watered seedlings (Fig. 1e, Fig. 2e; Bernacchi et al., 2002; Birami et al., 2020), while in the combined heat and drought treatment, *A_net_* decreased strongly with stress progression and declining *ψ*_Needle_ (Fig. 6a), concurrently with a heavy decline in *g*_s_ (Fig. S4; Ruehr et al., 2016; Birami et al., 2018). Under optimal water supply and heat, *g*_s_ also decreased (Fig. S3), which resulted in a close maintenance of *ψ*_Needle_ in isohydric pines. *A_net_* was modulated by a strong diurnal pattern in both stress treatments as apparent in larger C uptake during the cooler morning hours, directly after irrigation (Bauweraerts et al., 2013; Drake et al., 2018). The *A*_net_ decline was accompanied by a decrease in F’_v_/F’_m_ under heat stress in well-watered (Ameye et al., 2012; Guha et al., 2018) and with a more severe reduction in drought-treated seedlings (Fig. 2b; Birami et al., 2018). Other chlorophyll fluorescence parameters, i.e. electron transfer rate, effective photosystem II quantum yield and coefficients of photochemical fluorescence quenching have been shown to reveal strong decreases under heat and drought stress (Ameye et al., 2012; Birami et al., 2018). F’_v_/F’_m_ serves to assess stress impacts of the photosynthetic apparatus (Netto et al., 2005). Its decrease can be seen as a protective mechanism, i.e. a photoprotective process, which eliminates excess excitation energy, thus inhibiting the formation of harmful free radicals (Murchie and Lawson, 2013). The observed fast recovery of F’_v_/F’_m_ in the heat treatment is in line with previous studies (Ameye et al., 2012; Birami et al., 2018; Guha et al., 2018), but the recovery rate in drought-heat seedlings was lower. In summary, we found *A*_net_ to increase quickly to c. 20% of control rates alongside increasing F’_v_/F’_m_. This indicates that neither single nor compound stress had substantially harmed the photosynthetic apparatus.

Regarding *R*_root_, we found at first an increase in well-watered heat treated seedlings with rising temperatures (Fig. 1g, Fig. S5, Fig. S6) as commonly observed (Burton et al., 2002; Jarvi and Burton, 2013; Birami et al., 2020). The subsequent decline at soil temperatures >34°C is in agreement with a study on Aleppo pine seedlings, where *R*_root_ peaked at 31–34°C (Birami et al., 2020). This might indicate a respiratory acclimation response as shown also in a study on sugar maple (Jarvi and Burton, 2013). In drought-heat seedlings, *R*_root_ decreased strongly with drought progression, but apparently showed no large temperature sensitivity (Fig. S5). Explanations for declining respiration might be downregulated growth, lower availability of C, and at the cellular level limitation of substrates or adenylate control (Atkin and Tjoelker, 2003; Dusenge et al., 2019; O’Leary et al., 2019; Birami et al., 2020). Further, decreased C transport to sink organs might play a role due to reduced water cycling (Ruehr et al., 2009; Blessing et al., 2015). Despite lower *R*_root_ under drought-heat, the whole-tree C balance turned negative, which was hence dominated by a strong drop in *A_net_* (Fig. 3a, Fig. S6; Zhao et al., 2013), with *A*_net_ and C uptake halting or turning negative at a *ψ*_Needle_ of c. −1.0 MPa (Fig. 6a-b).

Following heat release in well-watered seedlings, the full recovery of shoot gas exchange is in agreement with a fully functional stem xylem (Fig. 4a-b). In contrast, delayed *A*_net_ recovery following drought-heat stress might be related to persistent reductions of *K*_Leaf_ and *K*_S_ compared to control seedlings (Brodribb and Cochard, 2009; Skelton et al., 2017; Rehschuh et al., 2020). *R*_root_ increased slowly during recovery from drought-heat stress, but surpassed control values at the end of the 18d-recovery-period (Fig. 1g). This reveals the enhancement of belowground repair mechanisms and root development to reestablish a fully functional root system (Davidson et al., 2006; Hagedorn et al., 2016), most likely to prepare for further stress periods.

Overall, we show that under heat stress, the C metabolism was affected moderately in well-watered and strongly in drought-treated seedlings. The decline of F’_v_/F’_m_ in both treatments indicates a protective mechanism of the photosynthetic apparatus, and its full recovery points towards no persistent photosynthetic damage. Alongside, *A*_net_ and respiration rates recovered almost fully, indicating a quick restoration of the C metabolism. This largely confirms our second hypothesis.

### Coupling of net C uptake and stem diameter change during stress and recovery

Stress-induced changes in water cycling and net C uptake simultaneously affected stem diameter change (Fig. 3). During the entire stress period, little stem growth appeared in drought-heat treated seedlings as cambium activity decreases at high xylem tension and low turgor (Abe et al., 2003; Balducci et al., 2013, 2016; Deslauriers et al., 2014), in agreement with the negative C balance (Fig. 3b). Further, stem shrinkage can be related to increased tree water deficit in phloem and xylem tissues. Under heat stress, the positive but reduced daily net C balance translated into slightly reduced stem growth as reported previously to occur under heatwaves (Bauweraerts et al., 2014; Ruehr et al., 2016). Following heat stress release, larger stem growth rates compared to control in well-watered seedlings were supported by a slight overcompensation of *A*_net_ and the associated higher net C balance (Fig. 1e, Fig. 3), reflecting stress compensatory responses (Balducci et al., 2016; Ruehr et al., 2016). Interestingly, in drought-heat seedlings, the C balance also recovered fast and reached control rates 2 d after rewetting. As *A*_net_ showed a tendency to remain below control rates while *R*_root_ recovered (on a tissue basis), this could be due to a relatively lower sink demand, as root biomass and the root:shoot ratio tended to be on average lower in drought-heat seedlings. The fast recovery of net C uptake suggests that the seedlings rebalanced C uptake and release, appearing largely independent of the reduced recovery of hydraulic conductance. We suggest that persistently lower *K*_S_ and *K*_Leaf_ in drought-heat seedlings did not translate into limitations of *A*_net_ and *E* at the generally low VPD conditions during the recovery period. During low VPD, the hydraulic system is operating far away from its maximum capacity, hence relatively small reductions in conductance as observed here have little effect on the water transport system. However, if another heat event occurred before the hydraulic system was fully restored, the persisting reductions of *K*_S_ and *K*_Leaf_ should limit *E* more strongly, thus reducing evaporative cooling and increasing thermal stress. Therefore, environmental conditions during recovery are an important factor to consider when drawing conclusions on the underlying mechanisms of stress legacy. In agreement, larger stem growth rates in drought-heat seedlings compared to control post-stress support a preference for the restoration of maximum hydraulic conductivity over the long-term (Brodribb and Cochard, 2009; Balducci et al., 2013).

Opposing to our third hypothesis, this indicates a fast regulation of the tree net C balance, appearing independent of hydraulic impairments under the recovery conditions applied here. However, enhanced stem growth post drought-heat supports the repair of the hydraulic system to restore maximum *K*_S_ and thus increase resistance to future stress conditions.

## Conclusion

Our study demonstrated that heat in combination with drought amplified the impacts on the carbon and water balance of Scots pine seedlings compared to heat stress alone. Well-watered heat treated seedlings were able to mitigate temperature stress by sustained evaporative cooling, while *g*_s_ and *A*_net_ declined moderately. This was also reflected in post-stress dynamics, indicating a fast recovery following heat stress with a slight overcompensation of net C uptake. In contrast, we found a delayed recovery of *R*_root_ and *A*_net_ after drought-heat release, while the photosynthetic apparatus appeared undamaged albeit high temperatures. Nonetheless, the net C balance in drought-heat seedlings recovered within 2 d after stress release, suggesting that a new equilibrium between C uptake and release was established. This was largely independent of the much slower recovery of *K*_Leaf_ and *K*_S_, most likely because the water transport system operated far away from its maximum capacity during the recovery conditions. Stem growth, however, might have been upregulated in order to produce new conductive tissues to repair the impaired hydraulic system. While our study shows that Scots pine seedlings are able to survive leaf temperatures >45°C alongside drought, it also indicates that seedlings are vulnerable to subsequent stress periods as the integrity of the hydraulic system, including supporting biomass, were not fully restored.

## Acknowledgements

We particularly thank Andreas Gast, Andrea Jakab, Benjamin Birami and Barbara Beikircher for technical and experimental support and advice. This study was supported by the German Research Foundation through its Emmy Noether Program (RU 1657/2-1) and by the German Federal Ministry of Education and Research (BMBF) through the Helmholtz Association and its research program ATMO.

## Author contributions

RR and NKR designed the study. RR conducted the experiment and analyzed the data with support from NKR. RR wrote the original draft with reviewing and editing from NKR.

## Notes

### Competing Interest Statement

The authors have declared no competing interest.

