## Supplemental material for "Unrevealing water and carbon relations during and after heat and hot drought stress in *Pinus sylvestris*"

#### Supplemental figures

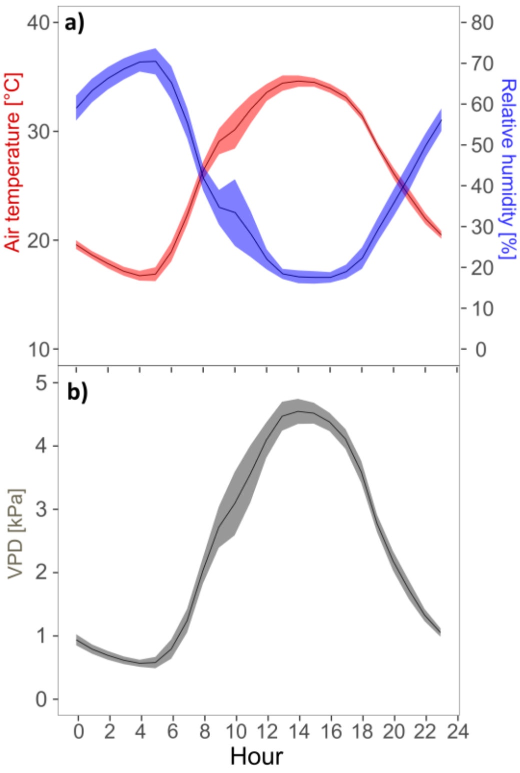


**Fig. S1: Diurnal progression of a) air temperature and relative humidity and b) vapor pressure deficit (VPD) averaged per hour for a heat period in August 2003 (07. - 13.08.) in Weißenburg- Emetzheim, Franconia, Germany. Around this place, Scots pine forest dieback has been reported following severe climate events. Data is from the meteorological station of the *German Meteorological Service*. Shown are hourly averages and shaded areas are ±SE.**

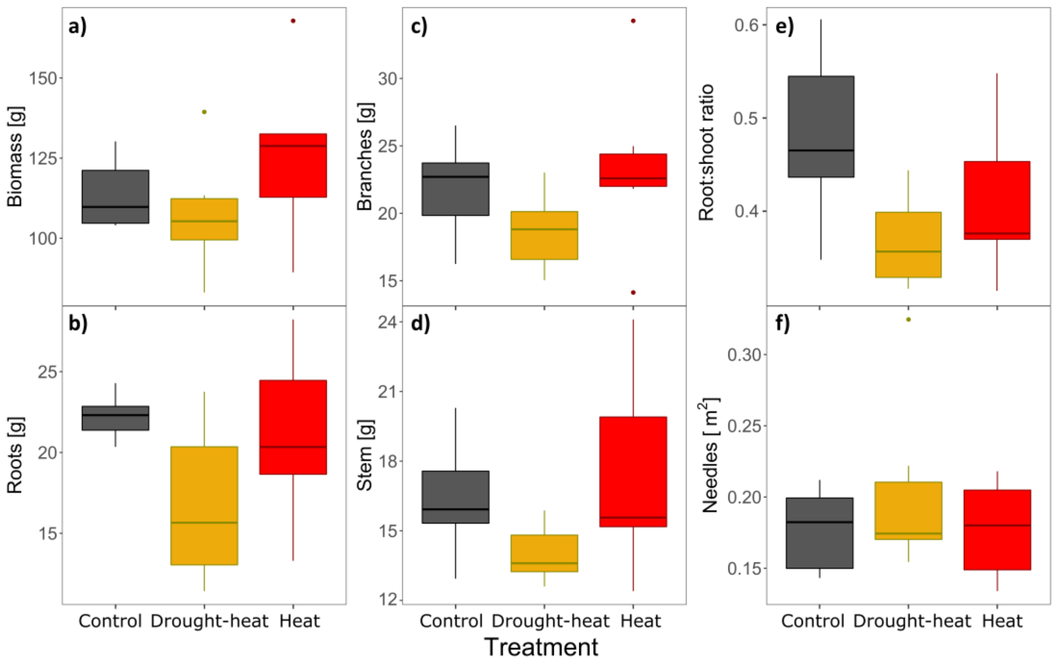


**Fig. S2: Biomass, root:shoot ratio and needle area of 3-year-old *Pinus sylvestris* seedlings in the tree chambers (n=6 per treatment). Dry weight of a) total tree biomass, b) roots, c) branches and d) stems, as well as e) root:shoot ratio, and f) needle area were determined at the end of the experiment. No significant difference between treatments was observed (Kruskal-Wallis test).**

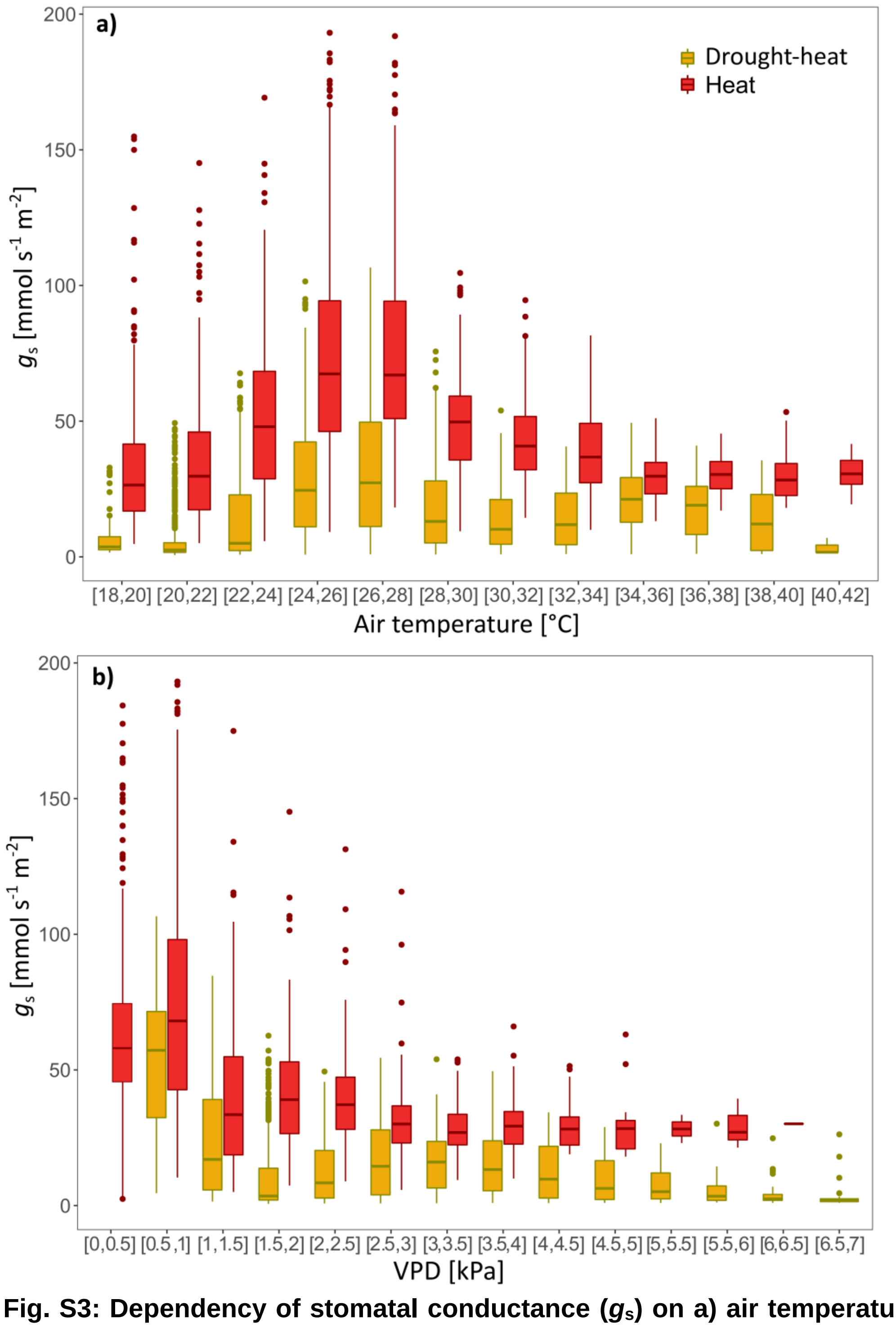


**Fig. S3: Dependency of stomatal conductance ($g_s$) on a) air temperature and b) vapor pressure deficit (VPD) for the stress treatments (n=6). Data are bin-averaged in temperature classes of 2°C and VPD classes of 0.5 kPa.**

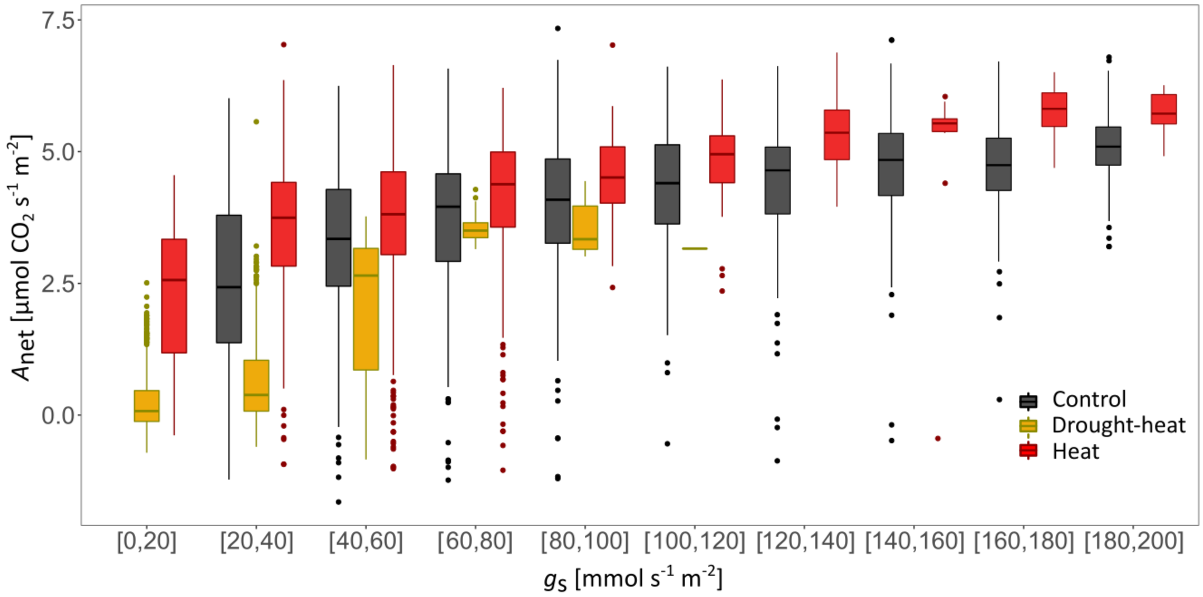


**Fig. S4: Dependency of net assimilation ($A_{net}$) on stomatal conductance ($g_s$) per treatment (n=6) during stress and control conditions. Data are bin-averaged in $g_s$ classes of 20 mmol $s^{-1}$ $m^{-2}$ (PAR>100).**

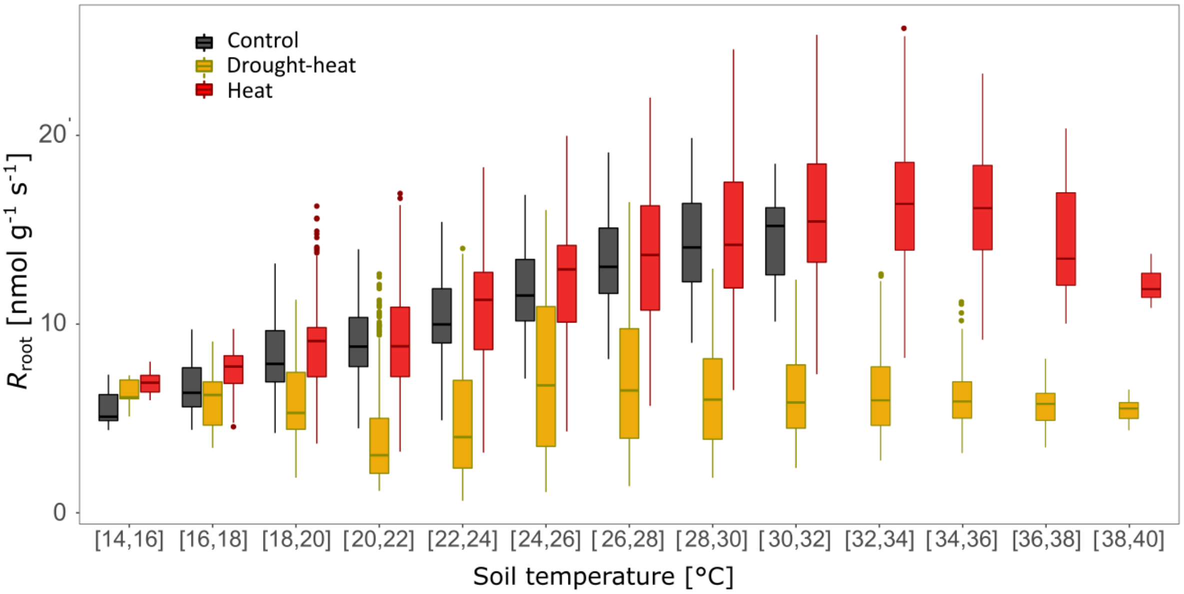


**Fig. S5: Dependency of root respiration ($R_{root}$) on soil temperature per treatment (n=6) during stress and control conditions. Data are bin-averaged in soil temperature classes of 2 °C.**

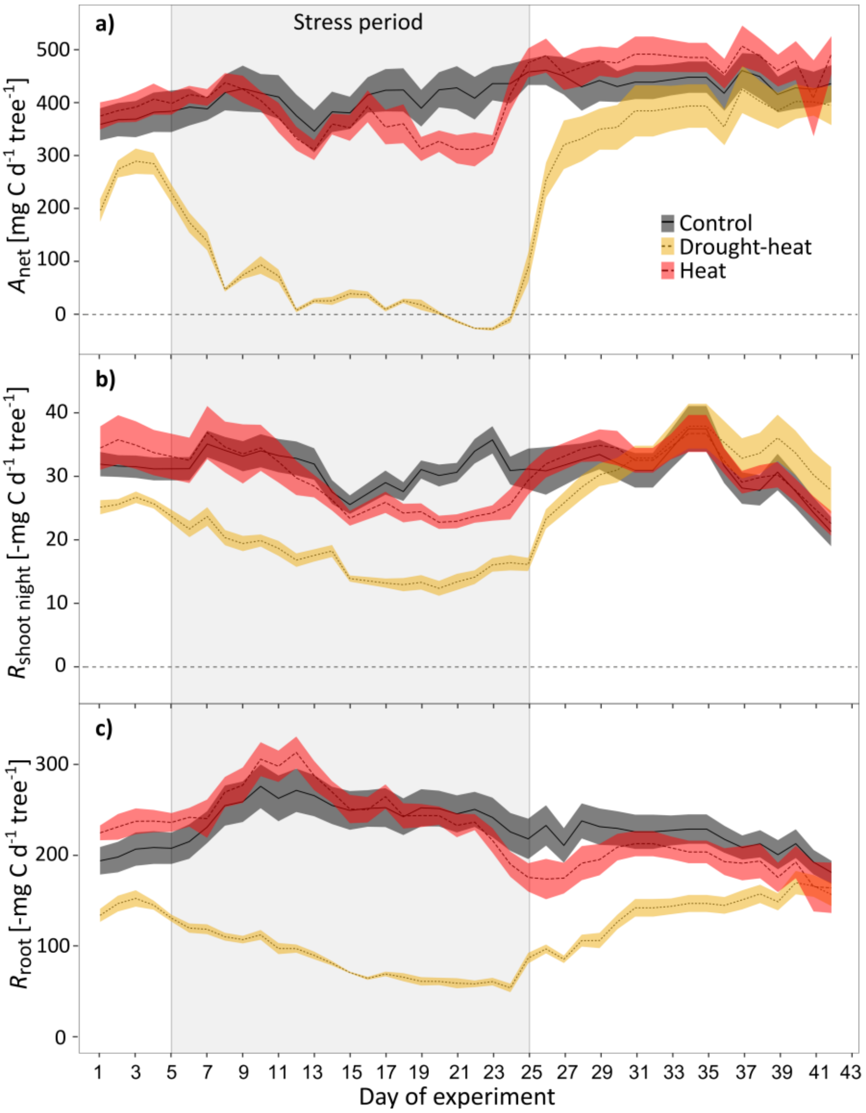


**Fig. S6: Dynamics of daily-averaged a) net canopy assimilation ($A_{net}$), b) shoot dark respiration ($R_{shoot\ night}$), and c) root respiration ($R_{root}$) of Scots pine seedlings. Data are treatment averages and shaded areas show ±SE (n=6). The gray boxes represent the stress period.**

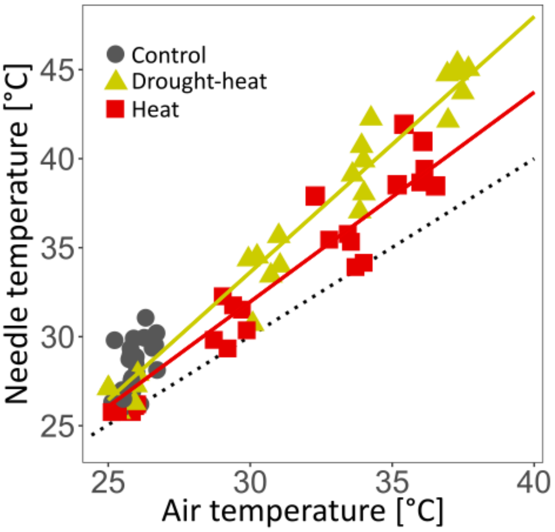


**Fig. S7: Leaf temperature was strongly correlated with air temperature for the heat ($r^2$= 0.93, p<0.05) and drought-heat treatment ($r^2$= 0.96, p<0.05), with well-watered seedlings (heat treatment) showing a higher capacity for leaf cooling. The dashed line shows the 1:1 relationship.**

### Supplemental tables

**Table S1: Functions for curves fitted to relationships of $A_{net}$, F´$_v$/F´$_m$ and $K_{Leaf}$ with leaf temperature for the heat and drought-heat treatment.**

| Correlation of max. leaf temperature with: | Heat | Drought-heat |
|---|---|---|
| $A_{net}$ | f(x)= -0.005 $x^2$ + 0.17 x +4.93 | f(x)= 0.015 $x^2$ -1.38 x + 30.67 |
| F´$_v$/F´$_m$ | f(x)= -0.001 $x^2$ + 0.08 x -0.96 | f(x)= -0.002 $x^2$ +0.10 x -1.32 |
| $K_{Leaf}$ | f(x)= 0.01 $x^2$ -0.74 x +15.68 | f(x)= 0.005 $x^2$ -0.42 x +9.86 |

**Table S2: Results of the Tukey post-hoc test of overall treatment comparisons per time period derived by linear-mixed effects models (lme) for continuous measurements, i.e. for net assimilation ($A_{net}$; daytime) and shoot night respiration ($R_{shoot\ night}$; nighttime), transpiration ($E$), root respiration ($R_{root}$) and net C uptake. Shown are p-values for all treatment comparisons for day and nighttime (diurnal measurements) and for the entire day for net C uptake. Significant p-values (P<0.05) are highlighted in bold. Tested periods include adjustment (26°C, day 1-4), temperature increments (35°C, day 10-16; 38°C, day 17-19; and 40°C, day 20-23), the initial (26°C, day 28-30) and final recovery period (26°C, day 40-42). DH= Drought-heat.**

| Parameter | Treatment | | Adjustment (daytime) | Adjustment (nighttime) | 35°C period (daytime) | 35°C period (nighttime) | 38°C period (daytime) | 38°C period (nighttime) | 40°C period (daytime) | 40°C period (nighttime) | Initial recovery (daytime) | Initial recovery (nighttime) | Final recovery (daytime) | Final recovery (nighttime) |
|---|---|---|---|---|---|---|---|---|---|---|---|---|---|---|
| $A_{net}$ / $R_{shoot\ night}$ [µmol m$^{-2}$ s$^{-1}$] | Control - | Heat | 0.93 | 0.99 | 0.99 | 0.99 | 0.83 | 0.83 | **0.0063** | 0.99 | 0.83 | 0.99 | 0.55 | 0.99 |
| | Control - | DH | **0.0014** | 0.97 | **<.0001** | **<.0001** | **<.0001** | **<.0001** | **<.0001** | **0.0096** | **0.0183** | 0.98 | 0.08 | 0.98 |
| | Heat - | DH | **0.0001** | 0.93 | **<.0001** | **<.0001** | **<.0001** | **<.0001** | **<.0001** | **0.0375** | **0.0009** | 0.91 | **0.0013** | 0.8 |
| *E* [mm m$^{-2}$ s$^{-1}$] | Control - | Heat | 0.99 | 0.96 | 0.99 | 0.99 | 0.974 | 1 | 0.89 | 0.43 | 0.71 | 0.99 | 0.94 | 0.94 |
| | Control - | DH | **0.0031** | **0.0433** | 0.058 | 0.058 | **0.0394** | **0.0362** | **0.0001** | **<.0001** | 0.95 | 0.43 | 0.87 | 0.87 |
| | Heat - | DH | **0.002** | **0.0087** | **0.024** | **0.024** | **0.0097** | **0.0351** | **<.0001** | **<.0001** | 0.25 | 0.26 | 0.42 | 0.42 |
| $R_{root}$ [µmol g$^{-1}$ s$^{-1}$] | Control - | Heat | 0.81 | 0.97 | 0.25 | 0.99 | 0.41 | 0.98 | 0.9 | 0.84 | 0.85 | 0.98 | 0.89 | 0.99 |
| | Control - | DH | 1 | 0.91 | **0.0014** | **0.0015** | **<.0001** | **0.0002** | **<.0001** | **0.0001** | **0.0168** | 0.07 | 0.87 | 0.86 |
| | Heat - | DH | 0.79 | 0.56 | **<.0001** | **0.0007** | **<.0001** | **0.0005** | **<.0001** | **0.0006** | 0.13 | 0.22 | 0.33 | 0.72 |
| Net C uptake [mg d$^{-1}$ tree$^{-1}$] | Control - | Heat | 1 | | 0.99 | | 0.99 | | 0.95 | | 0.95 | | 0.84 | |
| | Control - | DH | 0.07 | | **0.0006** | | **<.0001** | | **<.0001** | | 0.18 | | 1 | |
| | Heat - | DH | 0.08 | | **0.0097** | | **0.0006** | | **0.0001** | | **0.0061** | | 0.95 | |
